# Ancient dolphin genomes reveal rapid repeated adaptation to coastal waters

**DOI:** 10.1101/2022.11.03.515020

**Authors:** Marie Louis, Petra Korlević, Milaja Nykänen, Frederick Archer, Simon Berrow, Andrew Brownlow, Eline D. Lorenzen, Joanne O’Brien, Klaas Post, Fernando Racimo, Emer Rogan, Patricia E. Rosel, Mikkel H. S. Sinding, Henry van der Es, Nathan Wales, Michael C. Fontaine, Oscar Gaggiotti, Andrew D. Foote

**Author notes:** joint first author (Marie Louis, Petra Korlević, Milaja Nykänen). joint senior author (Michael C. Fontaine, Oscar Gaggiotti, Andrew D. Foote). Corresponding authors: Marie Louis, Andrew D. Foote.

## Abstract

Parallel evolution provides among the strongest evidence of the role of natural selection in shaping adaptation to the local environment. Yet, the chronology, mode and tempo of the process of parallel evolution remains broadly debated and discussed in the field of evolutionary biology. In this study, we harness the temporal resolution of paleogenomics to understand the tempo and independence of parallel coastal ecotype adaptation in common bottlenose dolphins (*Tursiops truncatus*). For this, we generated whole genome resequencing data from subfossil dolphins (8,610-5,626 years BP) originating from around the formation time of new coastal habitat and compared them with data from contemporary populations. Genomic data revealed a shift in genetic affinity, with the oldest ancient sample being closer to the pelagic populations, while the younger samples had intermediate ancestry that showed greater affinity with the local contemporary coastal populations. We found coastal-associated genotypes in the genome of our highest coverage ancient sample, SP1060, providing rare evidence of rapid adaptation from standing genetic variation. Lastly, using admixture graph analyses, we found a reticulate evolutionary history between pelagic and coastal populations. Ancestral gene flow from coastal populations was the probable source of standing genetic variation present in the pelagic populations that enabled rapid adaptation to newly emerged coastal habitat. The genetic response to past climatic warming provides an understanding of how bottlenose dolphins will respond to ongoing directional climate change and shifting coastlines.

## Introduction

Parallel genetic changes can arise from selection repeatedly acting upon standing genetic variation ^1,2^. However, how the timing and the independence of selection vary is still poorly understood for most natural study systems ^3,4^. For example, selection can act upon standing genetic variation in a shared ancestral population, or independently in multiple derived populations. Alternatively, adaptive standing genetic variation can be shared among derived populations through gene flow. Each of these non-mutually exclusive processes leaves a distinct genomic signature in the regions flanking the targets of selection ^3,4^.

In this study, we investigate the tempo and independence of parallel linked selection on standing genetic variation in coastal-adapted common bottlenose dolphins (*Tursiops truncatus*) derived from pelagic ancestors using contemporary and ancient (8,610-5,626 years BP) genomes. Coastal and pelagic ecotypes of bottlenose dolphins have recurrently formed in different regions of the world ^6–8^. Genetic variation has segregated under divergent selection between multiple coastal and pelagic ecotype pairs ^5^. In contrast, the same genetic variation is shared among widely distributed coastal populations ^5^. These coastal-associated variants are found mainly in ancient ancestry tracts in the genomes of coastal dolphins, and are present at low frequency as standing variation in pelagic populations ^5^. However, the mode and chronology of parallel linked selection on coastal-associated genetic variation in bottlenose dolphins remain unresolved. Here, we explore potential scenarios, incorporating ancient genomes dating from the estimated time of formation of local coastal populations ^8,9^.

Four subfossil samples dredged from the Southern North Sea bed in the eastern North Atlantic (ENA) were radiocarbon-dated to 5,979-5,626 calendar years before present (BP) for the youngest (SP1060, 95% CI) and 8,610-8,243 years BP for the oldest sample (NMR10326, Table S1, Figure 1a-b). These ages fall within the 95% CI range of the estimated split time of pelagic and coastal ecotypes in the ENA (47,800-4,300 years BP ^8^) and the emergence of coastal habitat in Northern European waters ^9^. This coastal habitat emerged when Doggerland was submerged by the North Sea around 12,000 to 6,000 years BP following post-glacial sea-level rise ^10^. During this warmer period of the early Holocene, subfossil evidence from the region confirms common bottlenose dolphin occurrence in the North Sea ^11,12^. It is thought this species entered the North Sea through the English Channel after it opened up following deglaciation ^11^.

**Figure 1.**
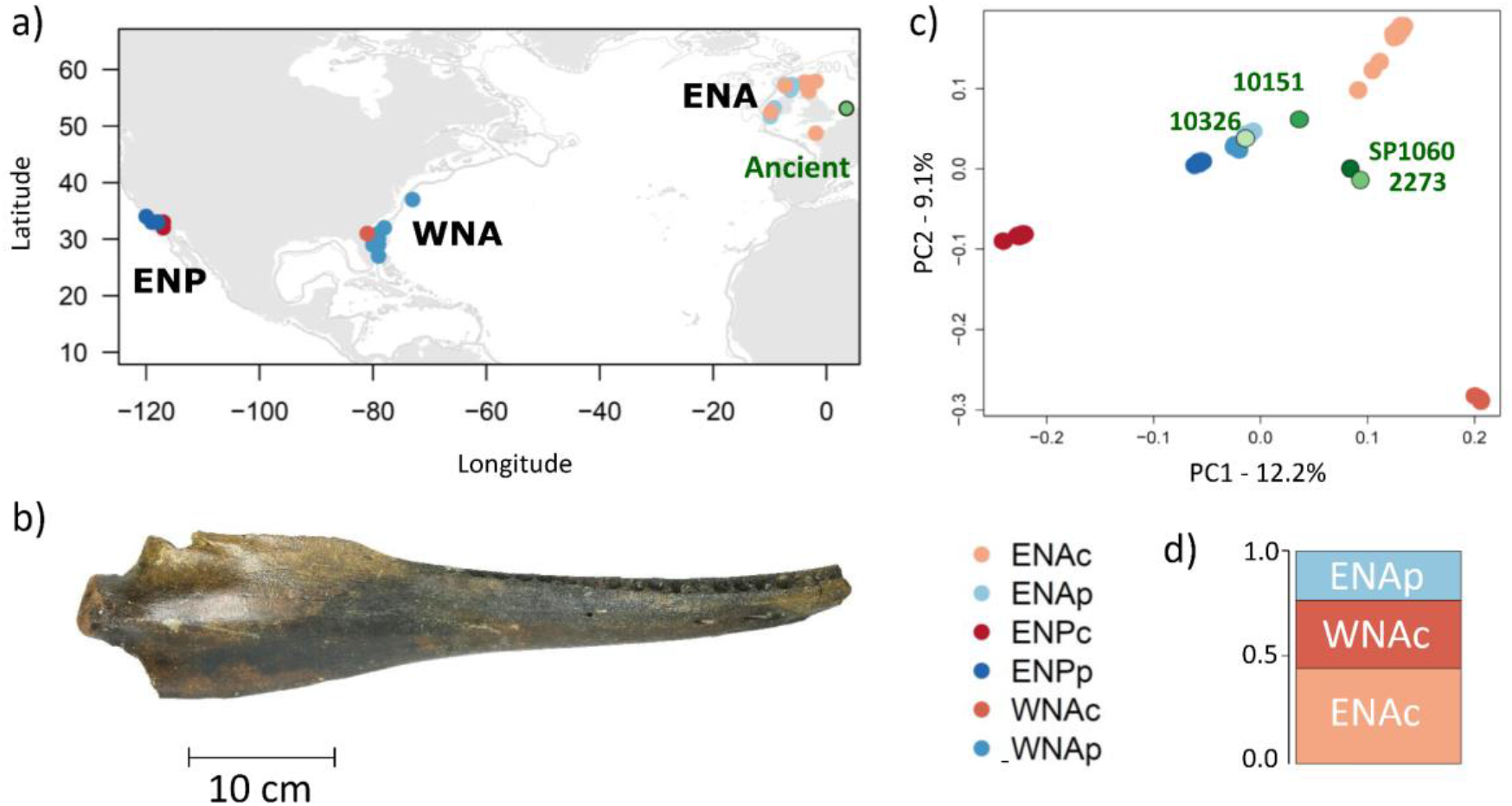
Sampling locations and ancestry of ancient and modern common bottlenose dolphin individuals. a) Map of sample locations of the four ancient (green) and 60 modern coastal (shades of red (c)) and pelagic (shades of blue(p)) common bottlenose dolphins in the eastern North Atlantic (ENAc and ENAp), western North Atlantic (WNAc and WNAp) and eastern North Pacific (ENPc and ENPp). b) Mandible of a 8,610-8,243 years BP bottlenose dolphin (sample NMR10326) included in this study. c) Principal component analysis of pseudo-haploid data from 60 modern and four projected ancient samples, mapped to the killer whale (*Orcinus orca*) reference genome to avoid reference bias and removing transitions, showing first and second principal components (PCs) based on 624,969 SNPs. The proportion of genetic variance captured by each component is indicated in the axes (see also Figures S4-S8). d) Ancestry proportions of sample SP1060 (5,979-5,626 years BP) identified using factorial analysis ^13^.

To understand the importance of these four early-to mid-Holocene ancient samples in the chronology of coastal adaptation, we first established their relationship to modern dolphins. We sequenced an ancient dolphin genome, SP1060, at 3x effective coverage (i.e. post-QC filtering, repeat masking, removing duplicates, base quality recalibration), to compare with a dataset of 60 modern genomes ^5^. We sequenced an additional three ancient dolphin genomes at ultra-low effective coverage (< 0.05x; Table S1) to verify whether SP1060 was representative of the genetic variation within the ancient population, rather than, for example, a rare admixed individual.

## Results and discussion

### Relationship of ancient genomes to modern dolphins

To explore the relationship between the ancient subfossils and modern individuals, we first looked at mitochondrial ancestry. The mitogenome sequence of the oldest ancient sample NMR10326 (Figure 1b) clusters with modern North Atlantic pelagic haplotypes, while the mitogenome sequences of the three other ancient samples clustered with modern Mediterranean and Black Sea samples in a Bayesian phylogenetic analysis (Figure S2). Then, on the nuclear data, we used principal component analysis ^14,15^ (PCA) and a factor analysis ^13^ (FA) method that can incorporate sample age, and therefore allows us to correct for temporal genetic drift when comparing populations ^13^. Given the differences in depth of coverage and the potential biases inherent in mapping and comparing ancient and modern genomes ^16^, we ran the multivariate analyses using several approaches: i) comparing pseudo-haploid genomes generated by sampling a random single base for all genomic positions and using PC projection of the ancient genomes on to the principal components segregating the modern genomes; ii) sampling a single read for each site with no projection but including sites covered in all modern and at least one ancient sample, and iii) comparing called genotypes for the modern individuals and SP1060 at sites with a minimum depth of 3x coverage, no missing data in SP1060 and both a projection and factorial analysis approaches. Due to the fragmented and damaged nature of ancient DNA, reads with non-reference alleles are less likely to be mapped than those with reference alleles, creating reference bias. To reduce reference bias, we ran those analyses with the genomic data mapped to the common bottlenose dolphin reference genome (GenBank: GCA_001922835.1), which is a coastal WNA individual, with relaxed parameters in BWA^17,18^, and again after re-mapping the data to the killer whale reference genome (Genbank GCA_000331955.2) ^19^. Using relaxed parameters or mapping to another species should help to include a better representation of alternate alleles. We also ran all analyses both including and excluding transitions, as C-T transitions are in excess at molecule ends of the SP1060 sequence reads, reflecting post-mortem deamination damage (see details in the Methods in the supplementary material). Additionally, we down-sampled one modern individual from the ENA coastal population to 0.03x to evaluate whether the lower coverage ancient samples could be pulled towards the middle of the PCAs due to large amounts of missing data. This ENA coastal individual clustered with the other ENA coastal individuals after downsampling in the PC projection (Figure S3) and single read sampling approaches (results not shown), indicating differences in coverage are not responsible for the observed patterns of variation.

As per Louis et al. (2021) ^5^, the major axis of differentiation is between the Pacific and the Atlantic populations (PC 1, Figure 1c). Independent genetic drift in each coastal population drives this pattern, while pelagic populations from both oceanic regions cluster in the centre of the PCA. Modern coastal samples from other locations in the ENA (northern France, Ireland and West Scotland) form a cline from the pelagic populations to the East Scotland samples. This could be consistent with a northwards range expansion of ENA coastal populations ^9^. The second axis of differentiation separated the two Atlantic coastal populations.

The ancient samples cluster with the pelagic samples along PC 2 (Figure 1c). Along PC 1, we observe affinity towards the two North Atlantic coastal populations for SP1060 and NMR2273, and to a lesser extent for NMR10151. We observe similar results in all analyses, regardless of filtering and mapping strategy (Figure 1c, Figures S4-S8). However, we note some reference bias for the ultra-low coverage samples (Figure S6). The oldest genome (NMR10326, 8,610-8,243 years BP) clusters with the pelagic populations when mapped to the killer whale reference genome (Figure 1c), while it does not do so when mapping to the bottlenose dolphin reference genome (Figure S6), likely due to reference bias associated with ancient DNA giving NMR10326 closer affinity to the WNA coastal population where the reference genome is from.

Having established that SP1060 is broadly representative of our ancient samples, we focus on SP1060 in the rest of our analyses. We estimated shared ancestry of SP1060 with the modern individuals using a factor analysis, which takes genetic drift into account ^13^. SP1060 shares the highest ancestry with the ENA coastal population (ENAc, 43%), followed by the WNA coastal (WNAc, 32%) and ENA pelagic (ENAp, 25%, Figure 1d) populations. Clearly, the inferred ancestry proportions do not represent an admixed ancestry composed of multiple modern populations, as SP1060 was alive at approximately the time of their divergence ^20^. Rather, they reflect ancestral genetic variation in SP1060, which later segregated in the different populations ^13,20^.

The PCA and ancestry results were further confirmed by the sharing of derived alleles using *D*-statistic tests ^21^ of the form *D*(H1,H2; SP1060,*Orca*). The ancient sample shares a significant excess of derived alleles with both ENA and WNA coastal populations, compared with all other in-group populations (Figure S10a). The value of the statistic *D*(ENAc,WNAc; SP1060,*Orca*) is significantly negative (Z-scores of -6.9 to -8.8), indicating that SP1060 is more closely related to the ENA coastal dolphins than to the WNA coastal dolphins. Accordingly, SP1060 shares a higher excess of derived alleles with the ENA coastal population than with the WNA coastal population, in statistics of the form *D*(coastal,pelagic; SP1060,*Orca*).

Having established the broad relationship of ancient samples to modern populations, in terms of shared ancestry, we next sought to reconstruct evolutionary history through time as an admixture graph ^22^. Testing across all possible histories, we find one admixture graph with no outlier *f*-statistics using qpBrute (Figure 2), which explores the space of all possible admixture graphs of a given maximum complexity, under a brute-force approach ^23,24^. Graphs were estimated using pseudo-haploid data mapped to the killer whale reference genome and using the killer whale as an outgroup. We find similar topologies when using called genotypes for the modern populations only and mapping to the bottlenose dolphin reference genome (Figure S11).

**Figure 2.**
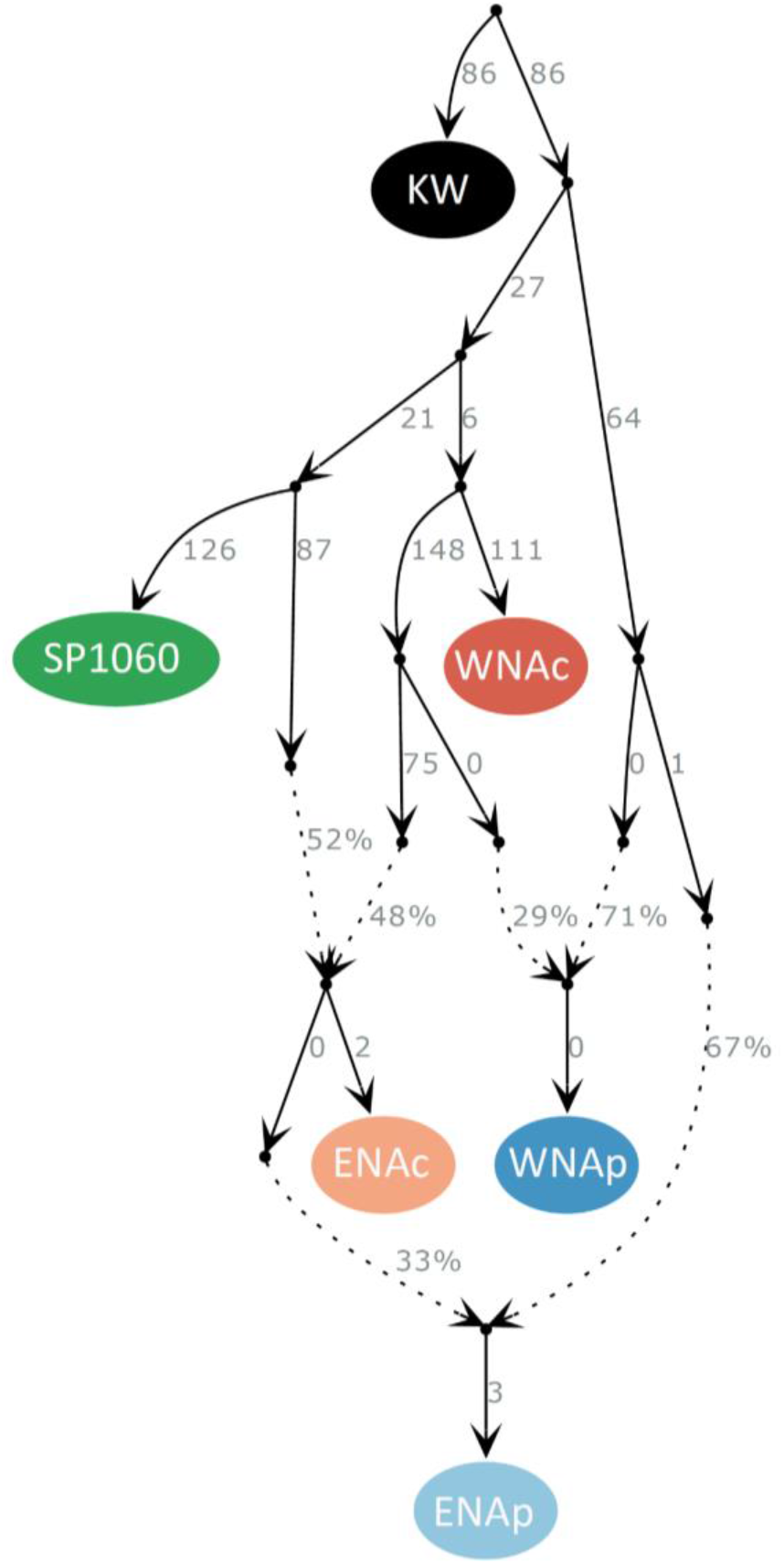
Evolutionary relationships between the ancient individual SP1060 and the North Atlantic modern bottlenose dolphin populations. Solid arrows indicate the relationships between populations/samples and the numbers at their right side the estimated genetic drift represented by the arrow. Populations include eastern North Atlantic coastal (ENAc) and pelagic (ENAp) populations, and western North Atlantic coastal (WNAc) and pelagic (WNAp) populations, and the outgroup is the killer whale (KW). Note that the drift value for SP1060 is inflated due to being a single and lower coverage-sample. Dashed lines are admixture edges and the arrows indicate the inferred direction of admixture, with the numbers reflecting the percentage of ancestry deriving from each lineage.

The best fitting graph reveals a basal split between the lineage that gave rise to the modern North Atlantic coastal populations and SP1060, and the lineage that gave rise to the majority of ancestry in modern pelagic populations (Figure 2). The WNA coastal, which has recently been described as its own species, *Tursiops erebennus ^25^*, and SP1060 are modelled as independent lineages. The ENA coastal is modelled as a clade whose ancestry is a mixture of the two ancestral groups leading to SP1060 and WNA coastal. Ancient sample SP1060 is therefore not a direct ancestor of the modern ENA coastal dolphins. The ancestry of both North Atlantic modern pelagic populations is modelled as an admixture of approximately 30% of the lineage giving rise to the coastal populations, and approximately 70% from a deeply divergent lineage. Thus, the model provides a useful visualisation of how coastal-associated alleles may have been reintroduced into pelagic populations. The branches leading to the pelagic populations have null or small drift values, consistent with pelagic populations having large ancestral effective population sizes or showing little genetic structure, and indicating minimal drift from a shared ancestral population, consistent with previous demographic inference ^5,26,27^.

### Patterns of selection to coastal habitat in the ancient individual

Coastal-associated genotypes at sites inferred as evolving under parallel linked selection in coastal populations ^5^ are also found in the ancient sample SP1060 (Figures 3a and b, S12). This observation informs us about the speed of selection. While we observe drift from the pelagic populations in the coastal modern individuals, SP1060 and the other ancient samples are genetically less diverged from the pelagic populations (Figure 1c). Yet, the genome of SP1060 shares the coastal associated variation inferred to have evolved under parallel linked selection in the coastal populations ^5^ (Figure 3a and b). The ancient genome shows excess heterozygosity as found in coastal individuals ^5^, despite its lower coverage (Figure S12). As SP1060 is dated to close or shortly after the colonisation time of coastal waters by bottlenose dolphins in the North Sea ^8,9^, this suggests selection occurred rapidly by acting upon standing genetic variation.

**Figure 3.**
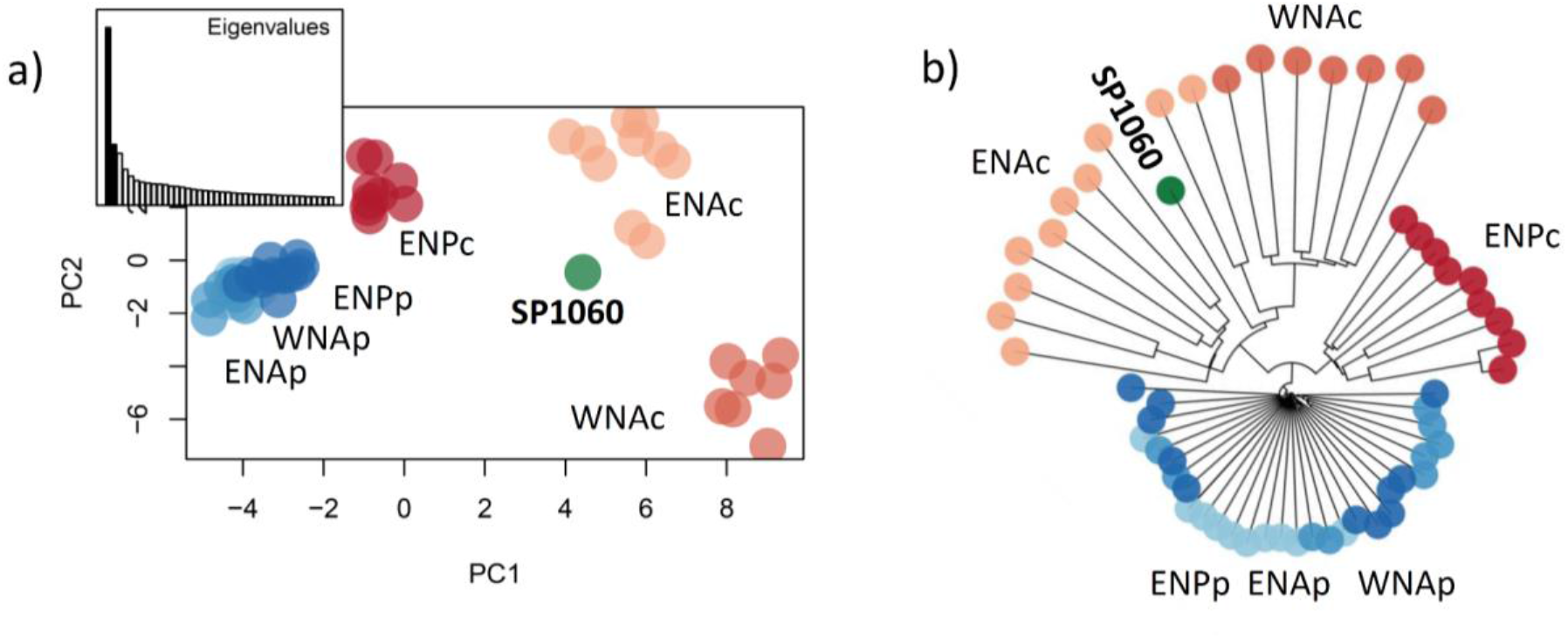
Patterns of genetic variation of the SNPs under repeated selection to coastal habitat in modern common bottlenose dolphin individuals and ancient individual SP1060. These include 2,122 SNPs with no missing data in SP1060 out of the 7,165 SNPs identified in Louis et al. (2021) ^5^, mapped to the bottlenose dolphin reference genome. Populations include coastal and pelagic ecotypes from the eastern North Atlantic (ENAc and ENAp), western North Atlantic (WNAc and WNAp) and eastern North Pacific (ENPc and ENPp). a) Principal component analysis and (b) Neighbour-joining distance tree showing the genetic structure of the common bottlenose dolphin samples for this particular SNP set.

Combining modern and ancient samples, we also provide insight on the independence of parallel linked selection in coastal bottlenose dolphins from standing genetic variation present in pelagic populations. Selection would be considered independent if it acted upon standing genetic variation in each derived population (scenario i) ^4^, that is along branches which are not shared among the two North Atlantic coastal populations, and SP1060 in Figure 2. It would not be independent if selection acted upon standing genetic variation in a shared ancestral population (scenario ii) ^4^, that is along branches shared among the coastal populations, or if adaptive standing genetic variation was shared post-selection among coastal populations through gene flow (scenario iii) ^4^.

To identify which of these three scenarios best fits our data, we compare patterns of variation (Figure S12) and the neighbour-joining tree (Figure 3b) obtained for our populations at the sites under parallel linked selection with the predictions under each scenario made by Lee and Coop 2019 ^4^. For the SNPs under parallel linked selection, the coastal individuals cluster by population, but with inter-individual variation ^5^. SP1060 clusters most closely with the ENA coastal population (Figure 3a), but diverges close to the basal node of the Atlantic coastal populations (Figure 3b). Such clustering of individuals by population is the pattern expected under scenario i), which describes an independent selection from standing genetic variation in each population, as it will generate only partially shared haplotypes. In contrast, scenarios ii) and iii) would have generated shared haplotypes among populations, and individuals from the North Atlantic coastal populations would be mixed in the tree. While the Lee and Coop approach is based on local trees, we based our analysis on different regions of the genome, thus assuming that the different regions fit the same selection scenarios. We compared the generated tree with ten trees generated using the same number of randomly sampled neutral SNPs. The trees do not show a clustering by ecotypes nor strong differences in branch length between coastal and pelagic individuals (Figure S13).

We observe long branches in the neighbour-joining tree for all the coastal individuals (Figure 3b). We previously identified these regions as representing old variants (0.6 to 2.3 million years); based upon the excess of mutations that had accumulated within them, compared with the genome-wide average ^5^. Given the findings of coastal-associated variants in the genome of SP1060, despite low drift from the pelagic population, we hypothesize that there was an abundance of coastal-associated standing genetic variation in the ancestral source population; for example, through a large influx of standing genetic variation from coastal populations in refugia, such as the Mediterranean Sea.

## Conclusion

The goal of this study was to disentangle the mode of parallel selection through direct observations of the chronology and genetic changes associated with the formation of coastal ecotype populations of bottlenose dolphins. Using subfossil samples pre-dating much of the drift experienced by coastal populations, we have disentangled neutral and selective processes. Overall, our results suggest that coastal variants represent balanced polymorphisms that were rapidly and repeatedly sieved from standing variation by ecological selection. We thereby provide rare direct evidence of rapid adaptation to newly emerged habitat from standing genetic variation.

## Methods

Detailed methods are provided in the supplementary material.

## Author contributions

ML, MN and ADF conceived the study; FA, SB, AB, ER, JOB, KP, PER, HvdE collected or curated samples/specimens, ML, PK, MN, MHSS, NW and ADF ran the laboratory work; MHSS and NW provided guidance in the ancient lab; ML and MN analysed the data; all authors interpreted the data; MCF, OEG, and ADF supervised the work, ML and ADF wrote the manuscript, with input from all co-authors. All authors read and approved the final manuscript.

## Acknowledgements

We especially thank Matthias Meyer for providing the SP1060 library, which allowed the generation of the ancient genome that was central to this study. We thank Ramon Fallon, Joseph Ward and Peter Thorpe from the St Andrews Bioinformatics Unit for help with bioinformatics. Bioinformatics and computational analyses were supported by the University of St Andrews Bioinformatics Unit which is funded by a Wellcome Trust ISSF award [grant 105621/Z/14/Z] and ran on cluster marvin and Crop Diversity HPC. We thank Kelly Roberston for the laboratory work for the modern samples from the USA and arranging shipment. We thank M. Thomas P. Gilbert for our use of the ancient DNA facilities in Copenhagen. We thank everyone involved in sample collection including Conor Ryan for sampling some of the stranded individuals in Ireland, the stranding networks in Ireland and Scotland and the Groupe d’Etude des Cétacés du Cotentin, and the NOAA Fisheries field crews and Simon Ingram for biopsy sample collection.

## Funding

Funding for ancient DNA labwork and sequencing was provided by the Total Foundation awarded to ML. Funding for sample collection in Ireland was provided by Science Foundation Ireland. Funding for labwork and visits to the University of Copenhagen was provided by the Systematics Research Fund, a Godfrey Hewitt mobility award from the European Society for Evolutionary Biology (ESEB), Lerner*-*Gray Grants for Marine Research. ML was supported by a Fyssen Fellowship, Total Foundation, the University of St Andrews and the Greenland Research Council. Modern sample DNA extractions were supported by People’s Trust for Endangered Species and the sequencing costs were supported by the European Union’s Horizon 2020 research and innovation program under the Marie Skłodowska-Curie grant agreement No. 663830 awarded to A.D.F., by the Total Foundation awarded to M.L., the University of Groningen awarded to M.C.F., and the Marine Alliance for Science and Technology for Scotland and The Russell Trust awarded to O.E.G. M.N. was funded by MASTS and the Crawford Hayes fund. A.D.F. was funded by Marie Skłodowska-Curie grant agreement No. 663830 and the European Research Council grant agreement No. ERC-COG-101045346.

## Notes

### Competing Interest Statement

The authors have declared no competing interest.

