## Supplementary material for "Ancient dolphin genomes reveal rapid repeated adaptation to coastal waters"

### Ancient dolphin genome reveals rapid repeated adaptation to coastal waters

Supplementary text

### MATERIAL and METHODS

#### Samples

Ancient sample collection and geological analyses

The four common bottlenose dolphin subfossil samples were dredged from the Southern part of the North Sea (i.e., Southern Bight and Smiths Knoll) by commercial trawlers, and were stored at the Natural History Museum of Rotterdam (Table S1). We collected bone powder in a sterile environment from the four specimens. Radiocarbon dating was performed in previous studies at the Klaus-Tschira-AMS facility for SP1060 [1] and at the University of Groningen for NMR2273 and NMR10326 [2]. For NMR10151, we performed radiocarbon dating at the University of Oxford. We re-calibrated the age of all the samples using the marine20 correction [3] in CALIB 8.2 [4] applying a ΔR (i.e. localised reservoir correction) of 8±38 as estimated for odontocete bones in Norway [5].

Modern sample collection

We used whole genome re-sequencing data from 57 bottlenose dolphins [6] and generated new data from three coastal individuals from the eastern North Atlantic. Two individuals were sampled using biopsy sampling in Northern France (Normandy) and Ireland (Shannon estuary). The third sample was collected from a stranded animal in West Scotland. Ecotype was identified previously using microsatellite markers [7]. Samples were either frozen or stored in 95% ethanol.

#### Laboratory work

##### Ancient DNA labwork

*Ancient sample SP1060*

Laboratory work was conducted in a dedicated ancient DNA facility at the Max Planck Institute for Evolutionary Anthropology in Leipzig for sample SP1060 (Table S1). The sample was treated with 0.5% bleach before extraction. Extraction and single-strand library preparation of the ancient sample are detailed in Korlevic et al 2015 [8]. The sample was sequenced on two lanes of HiSeq 2500 at the Danish National High-throughput Sequencing Centre of Copenhagen University.

*Ancient samples NMR2273, NMR10326 and NMR10151*

The three additional samples (Table S1) were processed at a dedicated ancient DNA facility at the Centre for GeoGenetics, University of Copenhagen. DNA was extracted from around 50 mg of bone powder, twice, both without and with a 1% bleach treatment. The samples were incubated overnight under motion at 55°C in 500 µl extraction buffer (0.45 M EDTA, 0.1 M UREA, 150 µg proteinase K). After centrifugation of the samples at 2,300 rpm for 5 min, the supernatant was collected, concentrated and purified using a Zymo-Spin V reservoir (Zymo Research Irvine, CA, USA) and Qiagen MinElute spin column (Qiagen, Inc., Valencia, CA, USA).

Double stranded libraries were built for these three samples following the blunt-end single tube library building (BEST) protocol [9]. Libraries were index amplified with the following thermocycling conditions: 2 minutes at 95°C, followed by 20 to 26 cycles of 30 s at 95°C, 30 s at 60°C, and 1 min50 s at 72°C, and a final, 10-min elongation step at 72°C. PCR products were purified using Qiagen MinElute spin columns (Qiagen, Hilden, Germany) following the manufacturer's instructions and eluted in 20 µl EB.

Two rounds of in-solution enrichment capture were performed on pooled libraries from different PCR reactions to increase complexity using a custom capture-enrichment kit produced by MYbaits (Mybaits Whole-Genome Enrichment, MYcroarray, Ann Arbor) [10]. RNA bait libraries were constructed by transcribing a fragmented modern high molecular weight bottlenose dolphin DNA (sample was from Foote et al. 2015 [11]). Ancient DNA libraries were hybridized to these RNA baits in a reaction containing adapter blockers for 24 hours. Streptavidin-coated magnetic beads were used to retain hybridized fragments and discard unbound DNA [12] following the manufacturer's instruction (online version 3.02 – July 2016). Captured libraries were amplified in two PCR reactions using KAPA HiFi polymerase (Kapa Biosystems) with the following thermocycling conditions: 2 minutes at 98°C, followed by 14 cycles of 20 s at 98°C, 30 s at 60°C, and 20 s at 72°C, and a final, 5-min elongation step at 72°C. Hybridization capture was performed a second time on the pool of PCR products. The double capture libraries were purified, quantified, pooled at equimolarity and sequenced on one lane of HiSeq 2500 with the 80 bp SE technology at the Danish National High-throughput Sequencing Centre of Copenhagen University.

##### Modern DNA labwork

DNA was extracted from epidermal tissue using a standard phenol:chloroform extraction method [13]. Genomic DNA was then sheared to an average size of ~500 bp using a Diagenode Bioruptor Pico sonication device. We built libraries on the DNA samples following the BEST protocol [9]. Libraries were pooled at equimolarity, together with samples for other projects and sequenced on an Illumina HiSeq 4000 with the 80 bp SE technology at the Danish National High-throughput Sequencing Centre of Copenhagen University.

#### Data processing

*Read trimming*

Illumina's CASAVA 1.8.2 software was used for the four ancient samples and the three new modern samples to convert of Illumina's *.bcl files to fastq and perform demultiplexing allowing for no mismatch in the 6-nucleotide indices used for barcoding. For the ancient samples, sequencing reads within the generated fastq files were processed with ADAPTER-REMOVAL v.2 [14] to trim residual adapter sequence contamination and to remove adapter dimer sequences as well as low-quality stretches at 3´ ends (i.e. consecutive stretches of N's and of bases with a quality score of 2 or lower). Sequence reads that were ≤30 bp following trimming were discarded. For the 57 modern samples from Louis et al. 2021 [6] and the three new modern samples generated during this study, sequencing reads were trimmed using Trimmomatic v.0.32 [15] using default parameters, and sequence reads shorter than 75 bp were discarded - see details in Louis et al. 2021 [6].

*Mitochondrial genomes*

The filtered reads were first mapped to a modified version of the bottlenose dolphin mitochondrial genome from Scotland (Genbank KF570351.1) [16] as per Morin et al. 2015 [17]. For the modern samples, we used BWA (v. 0.7.15) mem and default parameters [18]. For the ancient samples, we used BWA aln with the seed disabled (-l 1024) to include reads with post mortem damage at the read ends [19]. We also mapped the reads to two additional individuals, to ensure there was no reference bias, which can be an issue with ancient DNA, i.e. one individual from the Black Sea (Genbank KF570326.1) and one individual from the western North Atlantic (Genbank KF570378.1) [16]. We compared the sequences and checked they were the same. PCR duplicates were discarded using the rmdup function of samtools v. 1.2 [20,21] for the ancient samples and using picard-tools v. 2.1.0 [22] for the modern samples. A consensus sequence was constructed using the doFasta function in ANGSD v. 0.913 [23] setting the mapping quality to 25 and the phred score to 30. Nucleotide sites were changed to "N" if a single nucleotide did not represent >75% of the reads. All sequences were aligned with ClustalW in MEGA X [24] and visually inspected for indel and reading frames with 30 sequences from Nykänen et al. (2019) [25] and 75 sequences from Moura et al. (2013) [16].

*Nuclear genomes*

Reads that did not map to the mitochondrial genome were then extracted from the bam file and converted into a fastq file using samtools v. 1.2 and picard-tools v.2.1.0. For the modern samples, data were processed as described in Louis et al. (2021) [6]. In short, it involved the same steps as described below for the ancient samples, without the tweaks to decrease reference bias. For the ancient samples, these reads were then mapped to the reference bottlenose dolphin genome assembly (GenBank: GCA_001922835.1, NIST Tur_tru v1) using BWA (v. 0.7.15) aln (Li & Durbin 2009) with the seed disabled (-l 1024) and both default and adjusted parameters which were shown to reduce reference bias and include a better representation of alternate alleles (Martiniano et al. 2020) [26]. Those parameters were an edit distance (n) of 0.01 and a maximum number of gaps (o) of 2. To further evaluate reference bias, we also mapped the reads of the ancient and modern samples to the killer whale genome assembly (Genbank GCA_000331955.2) [11] using default parameters in BWA. Then, we kept only the mapped reads with a mapping quality of at least 25. We removed repeated regions as identified using RepeatMasker [27], regions of excessive coverage and mapping artifacts, and the sex chromosomes using a combination of bedtools and samtools (see details in Louis et al. 2021[6]). We also removed all the scaffolds which were shorter than 10 Mbp.

Endogenous content was inferred as the percentage of sequencing reads mapping to the reference genome after base quality filtering of q-score 30 and read duplicate removal (Table S2).

#### Data analyses

##### Assessing postmortem DNA damage and contamination

MapDamage 2.0 [28] was used to characterize post-mortem DNA damage and thereby check the authenticity of the data. Analyses revealed that the sequencing reads showed post-mortem damage characteristic of ancient DNA (Figure S1) including an excess of C/T transitions at both termini for SP1060 and an excess of C/T transitions at the 5' termini due to deamination and the G/A transition at the 3' termini for the three other samples on which double stranded libraries were built. Therefore, we used MapDamage to rescale base quality scores according to damage patterns when phred quality score >= 30. We then refiltered the bam files to keep only bases with a phred quality score >= 30. We ran all downstream analyses both including all sites and only including transversions. Results were the same, unless otherwise specified.

We detected no heterozygous sites in the mitochondrial genomes of the ancient samples, indicating an absence of contamination from present day DNA.

##### Mitochondrial DNA phylogeny

*Time-calibrated models using the mtDNA*

Mitochondrial genomes of *T. truncatus,* extracted from modern samples (n = 91), carbon-dated ancient subfossils (n = 4), or downloaded from GenBank (n = 48, see Table S3)[16] were first aligned in software MEGA X [1] with a *T. aduncus* reference mitogenome (KF570335.1) downloaded from GenBank. A topology tree in MrBayes [2,3], using 2,000,000 Markov Chain Monte Carlo (MCMC) samples and 25% burn-in, was then built with these sequences after the initial model selection for the best substitution scheme, Hasegawa-Kishino-Yano [4] with a proportion of invariable sites and gamma distributed rate variation among sites, in jModelTest2 [5,6]. The inspection of the resulting consensus tree revealed that one sample (sample 117696 collected from coastal eastern North Pacific) was very differentiated from the rest of the samples forming its own monophyletic branch. Therefore, the topology tree was also run without this sample using the same settings as before. The consensus trees were then inspected to find the two most evolutionarily distant *T. truncatus* haplotypes from both trees using the R-function 'cophenetic' from *ape* R package [7]. These sequences represented a sample obtained from coastal western North Atlantic (sample WNAC8) and the sample

117696; and sample WNAC8 and a sample originating from the eastern Mediterranean Sea (sample EMED6).

The estimation of time-calibrated phylogenies followed a two-step methodology by Morin et al. (2015) [17]. The purpose of the first step (delphinid tree model) was to estimate a calibration point (divergence of the two most evolutionarily distant *T. truncatus* haplotypes) for the second step (*T. truncatus* tree model).

*Delphinid phylogeny*

Thirteen protein coding genes of the mitochondrial genome from the two most divergent *T. truncatus* haplotypes (WNAC8 and 117696, or WNAC8 and EMED6) were aligned with the gene sequences of 19 delphinid species downloaded from GenBank (Table S3) [16,17,29–33]. The 'greedy' search in PartitionFinder (v1.1.0) [8] was used to find the best partitioning scheme for each gene, and two time-calibrated phylogenetic trees (one with and one without the sample 117696) were built in BEAST2 [9] with three different data partitions for nucleotide substitution models (Table S4). We used all codon positions and only the third codon positions, both with a single strict clock model based on the results from a previous study [10], and concatenated the individual gene trees into a single species trees to find the time to most recent common ancestor (TMRCA) between the two most divergent *T. truncatus* haplotypes. Two independent models for each scenario were run with 40,000,000 MCMC steps, 10% pre-burn-in, and a sampling frequency of 4,000. The divergence between Monodontidae and Delphinidae (mean=10.08 Myr, SD=1.413 [11]) was used to calibrate the root of the tree and a Calibrated Yule prior was used for the branching rate. The convergence of MCMC chains and the Effective Sample Size (ESS) values relating to the model parameters were checked in Tracer [12]. After verifying convergence, LogCombiner and TreeAnnotator [13] were used to combine and summarise the trees, respectively.

*T. truncatus phylogeny*

Thirteen protein coding gene regions were extracted from the 143 common bottlenose dolphin samples in order to estimate the coalescence times of different *T. truncatus* clades. Duplicate haplotypes were removed from this data set, thus eliminating one ancient sample, NMR2273 (duplicate with another ancient haplotype SP1060), and 52 modern samples. The best nucleotide partitioning scheme was determined for the remaining 90 haplotypes using

PartitionFinder (v1.1.0) [8] (Table S4). We used all codon positions and, as in Morin *et al.* (2015), only the third codon positions of the genes to minimise any possible effect of incomplete purifying selection on the coalescence times of the tree [14,15]. Time-calibrated phylogenies were built in BEAST2 [9] with two different data partitions for nucleotide substitution models (Table S4). We applied a common strict clock model for all of the genes, following Morin *et al.* (2015) [17] and Nykänen *et al.* (2019) [25]. For each tree built, two independent runs were performed with 100,000,000 MCMC steps, 10% pre-burn-in, and a sampling frequency of 10,000. Time to the most recent common ancestor (TMRCA) of all of the haplotypes, derived in the previous step when estimating the delphinid tree, was used to calibrate the root of the tree and a coalescent prior with constant population size was used for the tree branching rate. In addition, we applied tip calibrations with the carbon-dated sub-fossil samples. We ran altogether eight different scenarios for the trees, with and without the ancient sample NMR10151 which had low mitogenome coverage (2.3x) with 25% of the bases missing compared to the other subfossils, and with and without the modern sample 117696. The different tree scenarios are described in Table S5, and details of the priors can be found in the BEAST2 input .xml files. For all tree models, the convergence of chains and model performance was inspected in Tracer [12] after running each model twice, and LogCombiner and TreeAnnotator [13] were used to combine the log- and tree-files and summarise the trees, respectively. The resulting summary trees were drawn using FigTree v1.44 software (http://tree.bio.ed.ac.uk/).

#### Population structure

We used genotype likelihoods or pseudo-haploid data in ANGSD v.921 [23] to take into account differences in coverage between samples, where possible, or based our analyses on allele frequencies from called genotypes. We generated a vcf file including the modern individuals and SP1060 following the filtering steps as in Louis et al. 2021 [6]. In addition, in the vcf file, we kept only sites with a minimum individual coverage of 3x and sites with no missing data in the ancient sample.

*PC projection*

Note that all the analyses were run mapping to the common bottlenose dolphin reference genome with relaxed BWA parameters and to the killer whale reference genome to check there was no effect of reference bias. Results were the same, unless otherwise specified.

We performed PCA projections of the ancient genomes on the principal components segregating the modern genome with smartpca from the EIGENSOFT 7.2.0 package [34] using the option lsqproject. We run smartpca both on i) the modern samples and SP1060 and diploid genotype calls with a minimum individual depth filter of 3, and including the modern individuals, and ii) SP1060 and the three other ancient samples using pseudo-haploid genotypes generated in ANGSD v 9.21.

We converted the vcf genotype file to the eigenstrat format using the utility vcf2eigenstrat.py from https://github.com/mathii/gdc/blob/master/vcf2eigenstrat.py.

For the pseudo-haploid genotype calls, we first used the following filters in ANGSD, including a MAF of 0.05 (minMinor 6) and data in 25% of the individuals (maxMis16): -dohaplocall 1 -doCounts 1 -minMapQ 25 -minQ 30 -C 50 -baq 1 -minMinor 6 -maxMis 16 -skipTriallelic 1 -remove_bads 1 -uniqueOnly 1 -ref ref.fa

We then used the utility haplotoPlink from ANGSD to transform the haplo file to Plink tped and tfam. We used Plink 1.9 to transform those into bed/bim/fam files which we converted into eigenstrat using convertf from eigensoft/v6.1.3. We used the options --allow-extra-chr --chr-set 85 (killer whale reference genome scaffolds >10Mbp) or 56 (bottlenose dolphin reference genome scaffolds >10Mbp) in Plink.

*tfa method*

We also run the tfa method, which takes drift into account [35] on the diploid genotypes only, including the modern individuals and SP1060 and sites with no missing data in SP1060, mapped to the bottlenose dolphin reference genome with relaxed parameters. We imputed missing values for the modern samples using the package LEA for the run with the highest likelihood for K=6 (K=6 was previously described as the number of populations [6]) using the "impute" function in the R package LEA [36]. We also adjusted the genotypic data for coverage, as described in the method using the function "coverage_adjust" of the tfa R package, which uses a latent factor regression model. We used the function choose_lambda() to decide the lambda value for which the effect of sample age was removed from the fifth factor of the tfa analysis, similar values were obtained when choosing smaller factor values. We ran the tfa for K=5 as the two Atlantic populations are very closely related. We also estimated ancestry coefficients for SP1060 including iteratively all populations. SP1060 did not show any ancestry relationship with the Pacific populations. Therefore, we estimated the ancestry coefficients for SP1060 including ENAc, WNAc and ENAp populations (including WNAp instead of ENAp gave the same results, likely due to the two populations being closely related).

*ANGSD single-read sampling approach*

We also ran the single read PCA method in ANGSD v.0.921, which involves random sampling of a single read for each sample at each site. In this analysis, the ancient samples were included in the PC computations and not projected onto PCs of modern samples. This provides a quality control measure, such as if the ancient samples presented sequencing or sequence data processing errors, they would appear as outliers in the PCA in comparison with the modern samples. We ran the analyses, with and without the transitions, and on the data mapped to the bottlenose dolphin reference genome and the killer whale reference genome, on i) the 60 modern individuals and SP1060, allowing no missing data, ii) the 60 modern individuals, SP1060 and NMR10151 allowing no missing data, iii) the 60 modern individuals, SP1060, NMR2273 and NMR10326 allowing for missing data in one individual, and iv) the 60 modern individuals and all four ancient individuals allowing for missing data in two individuals. As results were consistent, we only present those including all the samples.

We used the following options in ANGSD: samples.bamlist -nThreads 9 -doIBS 1 -doCounts 1 -doMajorMinor 1 -minFreq 0.05 -minInd 61 -maxMis 2 -rmTrans 1 -output01 0 -makeMatrix 1 -doCov 1 -minMapQ 25 -minQ 30 -out out_rmTrans -GL 1 -skipTriallelic 1 -remove_bads 1 -uniqueOnly 1

#### Evolutionary relationships

*D-statistics*

*D*-statistics were used to assess the relationships of ancient sample SP1060 to the modern populations. *D*-statistics were first performed on all 15 possible combinations including two modern samples and the ancient sample, with the killer whale, *Orcinus orca*, as the outgroup. The *D*-statistic describes an excess of shared derived alleles between taxa which could be the result of introgression or ancestral population structure. It thus allows to detect departure from ‘tree-ness’ of a given topology [37–39]. We considered that H1 and H2 are two modern dolphin populations. We used the *D*-statistics to evaluate if the data are consistent with the null hypothesis that the tree (((H1,H2),SP1060),*Orca*) is correct and that there has been no gene flow between the ancient sample and neither H1 or H2. The definition of the *D*-statistics used here is the one of Durand *et al.* 2011.

D=(nABBA-nBABA)/(nABBA+nBABA) where nABBA is the number of sites where only H2 and H3 share a derived allele (ABBA sites) and nBABA is the number of sites where only and H1 and H3 share a derived allele (BABA sites). Under the null hypothesis that the given topology is the true topology, we expect an equal number of ABBA and BABA sites and thus *D*=0. A statistic differing significantly from 0 indicates either gene flow between one population within the in-group and H3, or that the tree is incorrect. We implemented the tests in ANGSD v 0.921, and sampled a single base at each position of the genome to remove bias caused by differences in sequencing depth and only considering sites covered in all individuals. *D*-statistics were calculated using one individual per population and the test was repeated three times using a different individual each time. We ran the analyses on the data mapped to the common bottlenose dolphin reference genome with relaxed BWA parameters and to the killer whale reference genome to check for reference bias, and both without and keeping the transitions, to make sure there was no impact from DNA damage patterns. We used the same filters as described above for the PCAs. The significance from deviation from 0 was assessed using a *Z*-score based on blocked jackknife estimates of the standard deviation of the *D*-statistics (block size was 5Mb which should be higher than the LD in the populations). This *Z*-score relies on the assumption that the *D*-statistics, under the null hypothesis, is normally distributed with mean 0 and a standard deviation equal to a standard deviation estimate computed using the “delete-m jackknife for unequal m” method described in Busing et al. 1999 [40]. An example of a command is: angsd -out ${1}_XchrmHWE_minIndKWcov -doAbbababa 1 -rmTrans 1 -blockSize 5000000 -enhance 1 -bam ${1}_XchrmHWE.filelist -doCounts 1 -useLast 1 -minMapQ 25 -minQ 30 -minInd 4 -maxMis 0 -remove_bads 1 -uniqueOnly 1

*Admixture graph analysis*

We next reconstructed phylogenetic relationships between SP1060 and the Atlantic modern populations (n=37) using an admixture graph analysis [37]. We excluded the Pacific populations from this analysis as there are many other populations in between the regions. We only included the already published ENAc individuals (n=10) and did not include the newly generated three individuals as they belong to different populations [7].

We used a heuristic search algorithm, qpBrute (https://github.com/ekirving/qpbrute) [41,42] to explore the space of all possible admixture graphs fitted to the four Atlantic populations using qpGraph. We built the admixture graph using two dataset: i) pseudo-haploid data including SP1060 and the modern Atlantic populations mapped to the killer whale reference as it minimizes reference bias, and ii) called genotypes for the modern Atlantic populations mapped to the bottlenose dolphin reference. We used these two datasets to check results were similar, irrespective of the data type (pseudo-haploid and called genotypes) and reference genome.

For the pseudo-haploid data, we used 580,589 SNPs and a killer whale genome as the outgroup. We kept the transitions as evolutionary relationships were similar with and without the transitions (Figure S9). We used the following command in ANGSD: angsd -b SP1060_Atlantic_KW_10MbpKW.bamlist -dohaplocall 1 -doCounts 1 -minMapQ 25 -minQ 30 -C 50 -baq 1 -minMinor 4 -maxMis 9 -skipTriallelic 1 -remove_bads 1 -uniqueOnly 1 -ref /storage/home/users/ml228/Genomics/NEA_BGI_data_ALL/NEA_BGI_clean_raw_data/Refs/unpla/unplaced.scaf.fa -out Atlantic_KW_SP1060_mapKW

We then converted the data to the eigenstrat format as described above for smartpca.

For the modern called genotype data, we used one Indo-Pacific bottlenose dolphin, *Tursiops aduncus*, individual to root the graph. We filtered the dataset as follows: a mapping quality of 30, a genotype quality of 20, genotype depth of at least 3x, less than 25% of missing data overall, genotype data in at least 5 individuals in each population, a MAF of 0.05, bi-allelic SNPs only, no missing data in the *T. aduncus* individual constituting the outgroup, scaffolds of at least 10 Mbp, and a distance of at least 5 Kb between SNPs. We obtained 223,950 SNPs. We also set the missing data to less than 10% and got similar results.

We ran qpBrute using the following parameters: outpop: NULL, useallsnps: YES, blgsize: 0.05 (5Mb which is the block size for Jackniffe), forcezmode: YES, lsqmode: YES, diag: .0001, bigiter: 6, hires: YES, lambdascale: 1, inbreed: YES (for the pseudo-haploid data only).
As we have only one individual, we did not attempt to use the allele frequency of the outgroup to normalise the weighting of each SNP in the ingroup and use output: NULL that is SNPs are flat weighted. We used the option inbreed:YES for the pseudo-haploid data. This allows for each pseudo-haploid to contribute to only one haplotype when applying the low low-sample-size correction for the *f2*-statistics. This does not work when including only a single pseudohaploid sample as we did with SP1060, but it should not be an issue as using the default option inbreed:NO gave similar results.

In qpBrute, leaf nodes were added to the graph using a stepwise addition order algorithm. At each step, insertion of a new node was tested at all branches of the graph, apart from the outgroup branch. All possible admixture combinations were tried where a node could not be added without producing *f4*-statistics outliers (i.e., $|Z| \geq 3$). The sub.graph was discarded when a node could not be inserted via these approaches. Where a node was successfully added, then the remaining nodes were recursively inserted into the graph.
For the pseudo-haploid dataset, the package tried all possible 120 starting graph orders. We found only one graph with no *f4* outliers among a total of 3,663 unique graphs.
For the called genotypes, the package tries all possible 24 starting graph orders and fitted 234 unique admixture graphs to our data set. We found three possible graphs with no *f4* outliers left, although two of them were mirror graphs. We computed the mean log likelihoods of the three models and their Bayes Factors using the MCMC algorithm implemented in the R package ADMIXTUREGRAPH [43] using the default chain settings implemented in qpBrute (two chains, each with two million iterations, five heated chains, a burn in of 50%, and no thinning). We assessed convergence of the chains using the output from the R package CODA [44], also generated in qpBrute. The Bayes Factors showed a non-significant support of the first model in comparison to the other “mirror” two, which have similar log likelihoods (Bayes Factor of 0.88). We increased the chains to four millions iterations with a burnin of 3 millions, but it did not help in discriminating between the three graphs.

### RESULTS

#### Mapping statistics

##### *Mitogenomes*

The *T. truncatus* mitogenome sequence has 16,391 sites. We called three nucleotides in NMR10151 and one nucleotide in NMR2273 as Ns as there was not a single nucleotide representing > 75% of the reads. There were no ambiguities in SP1060 and NMR10326. In the 61 modern samples, there was one ambiguous base (i.e. without a single nucleotide representing > 75% of the reads) in four of the samples (160321, IR33, S37, S41), which was changed to N. This could represent heteroplasmy.

Coverage of the mitogenomes for the ancient samples was as follows: SP1060: 160.4x, NMR2273: 9.9x, NMR10326: 27.6x and NMR10151: 2.3x.

Average mitochondrial genome coverage of the 57 modern samples from Louis et al. 2021 [6] was 1446.2x (SD=595.1) and the new four modern samples from this study was 103.4x (SD=16.4).

Proportion of missing bases (N) was 0.0001 in the 57 modern samples (122 bases over all samples), 0.006 in the three new modern genomes (3 bases overall all samples), 0.0002 for SP1060 (4 bases), 0.003 for NMR10326 (50 bases), 0.006 for NMR2273 (100 bases) and 0.25 for NMR10151 (4264 bases).

##### *Nuclear results statistics*

For SP1060, endogenous content was 28% when mapping to the bottlenose dolphin reference genome and 24.1% when mapping to the killer whale reference genome, and coverage was 3x (Table S2). Mapping statistics for all samples are detailed in Table S2.

#### Mitochondrial DNA phylogeny

##### *Delphinid phylogeny*

The average overall rate for substitutions per site per million years (clock rate) for the Delphinids in both phylogenies (with and without the sample 117699) was estimated as 6.670 $\times 10^{-3}$ (95% HPDI: $4.877 \times 10^{-3} - 8.835 \times 10^{-3}$), and the TMRCA of the two most divergent *T. truncatus* samples was estimated as 0.916 Mya (95% HPDI: 0.630 – 1.234 Mya) for the phylogeny without 117696 and as 1.174 Mya (95% HPDI: 0.814 – 1.567 Mya) for the tree including 117696. The Effective Sample Size (ESS) values for the different parameters in the combined runs were all >250, with most of them >3,000, indicating no autocorrelation between samples.

*T. truncatus phylogeny*

The ESS values for the model parameters in the *T. truncatus* tree models were all >1500 (in individual runs), indicating no sign of autocorrelation between samples and a good convergence of chains. The scenario with the highest posterior probabilities of deeper (older) nodes included all of the subfossil samples and the modern sample 117696. We will therefore present only the results from this model, however, it is important to note that all model scenarios (all codon and 3rd codon) placed the coastal NWA samples into a separate monophyletic clade, concordant with recent studies [16,17]. In addition, the placement of the subfossil samples was identical in all model scenarios (Figure S2).

The average clock rate for the all-codon bottlenose dolphin phylogenetic model was estimated as $7.026 \times 10^{-3}$ substitutions/site/Myr with 95% HPDI of $4.579 \times 10^{-3}$ – 9.949. The summary consensus tree consisting of *T. truncatus* samples indicates that the coastal WNA clade has the oldest coalescence time of 0.8 My (95% HPDI: 0.493 – 1.100 My) (Figure S2). A clade consisting of samples collected from the eastern North Pacific, however, has a slightly younger mean coalescence time of 0.704 My (95% HPDI: 0.442 – 0.984 My). All of the dated subfossil samples are placed in a clade that include mostly modern samples collected from pelagic North Atlantic, however, there are also three samples in this clade originating from the Mediterranean and Black Seas (Figure S2). As in a previous study [10], incomplete lineage sorting is evident in the coastal eastern North Atlantic sequences.

#### Population structure

*PC projection*

The position of SP1060 is consistently intermediate between the pelagic populations and the two North Atlantic coastal populations in all analyses on the first and second PC, that is when using called diploid genotypes (Figure S4), or pseudo-haploid calls (Figure 1C, Figure S4), removing (Figure 1C, Figure S6) or keeping transitions (Figure S4). The coastal and the pelagic samples separate on PC3, with the ancient sample being intermediate (Figure S5).

While the position of SP1060 remains relatively unchanged when mapping to the bottlenose dolphin (Figure S6) or the killer whale (Figure 1c) reference genomes, we note some reference bias when mapping the ultra-low coverage samples to the bottlenose dolphin reference genome (Figure S6). The position of the three samples is shifted towards being closer to the WNAc individuals, the reference genome being from a WNAc population, than when mapping to the killer whale reference genome (Figure 1c), in particular for the lowest coverage sample NMR10151.

*tfa method*

In the tfa analysis, factor 1 separates the Pacific and the Atlantic population, while factor 2 separates the coastal populations from the ENA and WNA. The tfa analysis also indicates that SP1060 is intermediate between the two Atlantic coastal populations and the pelagic populations (Figure S5).

*ANGSD single-read sampling approach*

Using the single read sampling method in ANGSD we got relatively similar PCA results than with smartpca. The position of SP1060 as intermediate between the two Atlantic coastal populations and the pelagic is unchanged (Figure S8). We find that the two younger samples (SP1060 and NMR10151) are closer to the coastal populations than the two older samples (NMR2273 and NMR10326, Figure S8).

#### Evolutionary relationships

*D-statistics*

The *D*-statistics results were consistent for the data mapped to the bottlenose dolphin reference genome (Figure S9) and the killer whale reference genome (Figure S10), with the transitions (Figures S9a, S10a) and removing the transitions (Figures S9b, S10b). We note some slight reference bias towards the WNAc when mapping to the WNAc bottlenose reference genome, in particular when including the transitions. The value of the statistic *D*(ENAc,WNAc; SP1060,*orca*) is significantly negative, indicating that SP1060 is more closely related to the ENAc dolphins than the WNAc dolphins. This pattern is the strongest when mapping to the killer whale genome, with and without transitions (Figure S10), and the lowest when mapping to the bottlenose dolphin reference and when keeping the transitions (Figure

S9b), highlighting the need to take reference bias and damage patterns into account. Similarly, ENAc shared a higher excess of derived alleles with SP1060 than WNAc, with the value of statistics of the form *D*(coastal,pelagic; SP1060,*orca*) being lower when the eastern Atlantic populations are included than when the western are. This pattern is the strongest when mapping to the killer whale reference genome.

*Admixture graph analyses*

We find similar graph topologies when using the pseudohaploid data mapped to the killer whale genome and including both SP1060 and the modern Atlantic populations (Figure 2) and when using called genotypes and the modern Atlantic populations only (Figure S11).

### Supplementary figures and tables

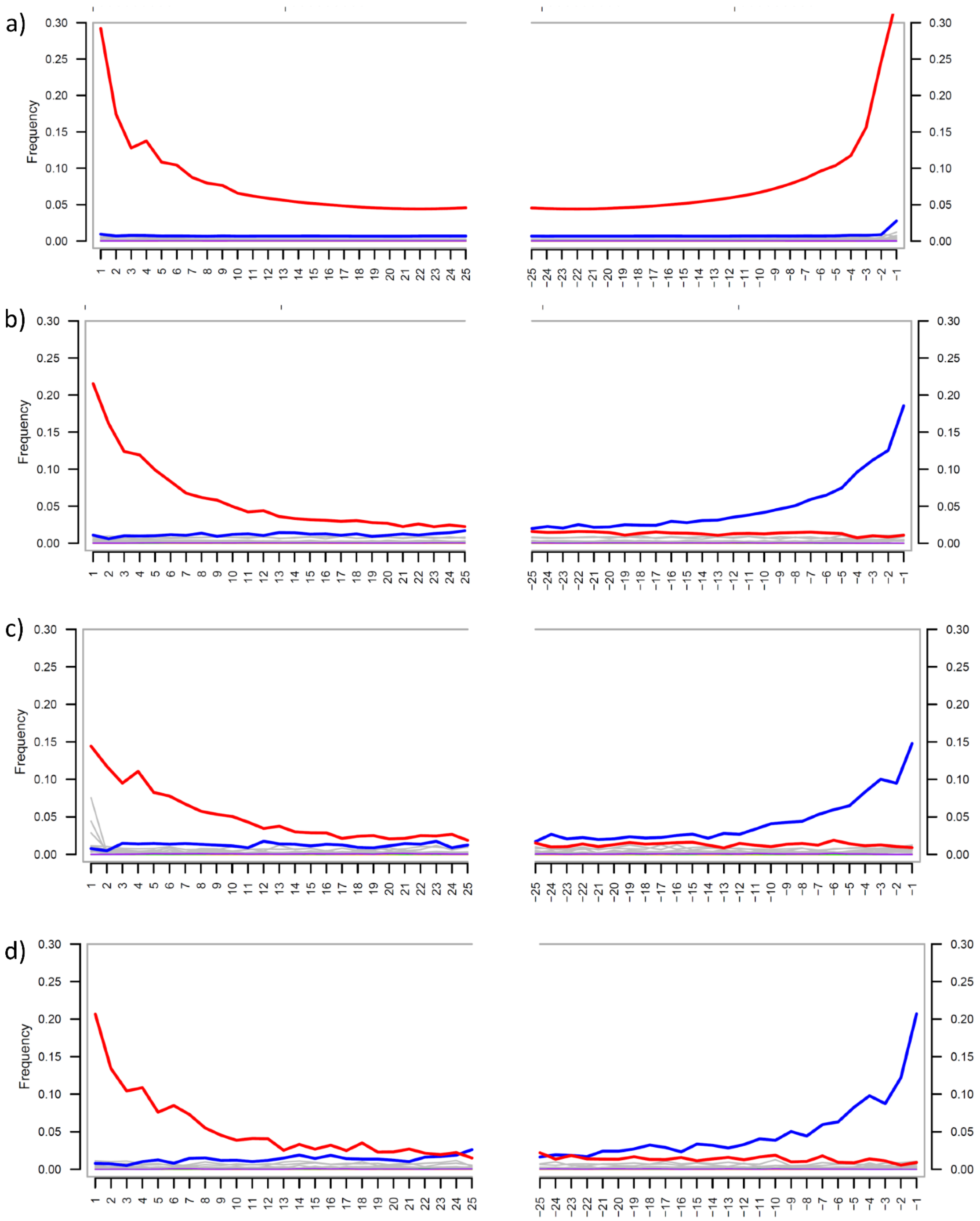

Figure S1. Misincorporation pattern plot obtained with MapDamage for the first and last 25 bases for a) SP1060, b) NMR10326, c) NMR2273 and d) NMR10151. It shows the percentage of sites containing a nucleotide change from the killer whale reference sequence along the DNA fragment with: red indicating C to T transitions, blue indicating G to A transitions. SP1060 doesn't show the complementary G->A excess due to being a single stranded library. Grey represents all other substitutions and purple insertions relative to the reference.

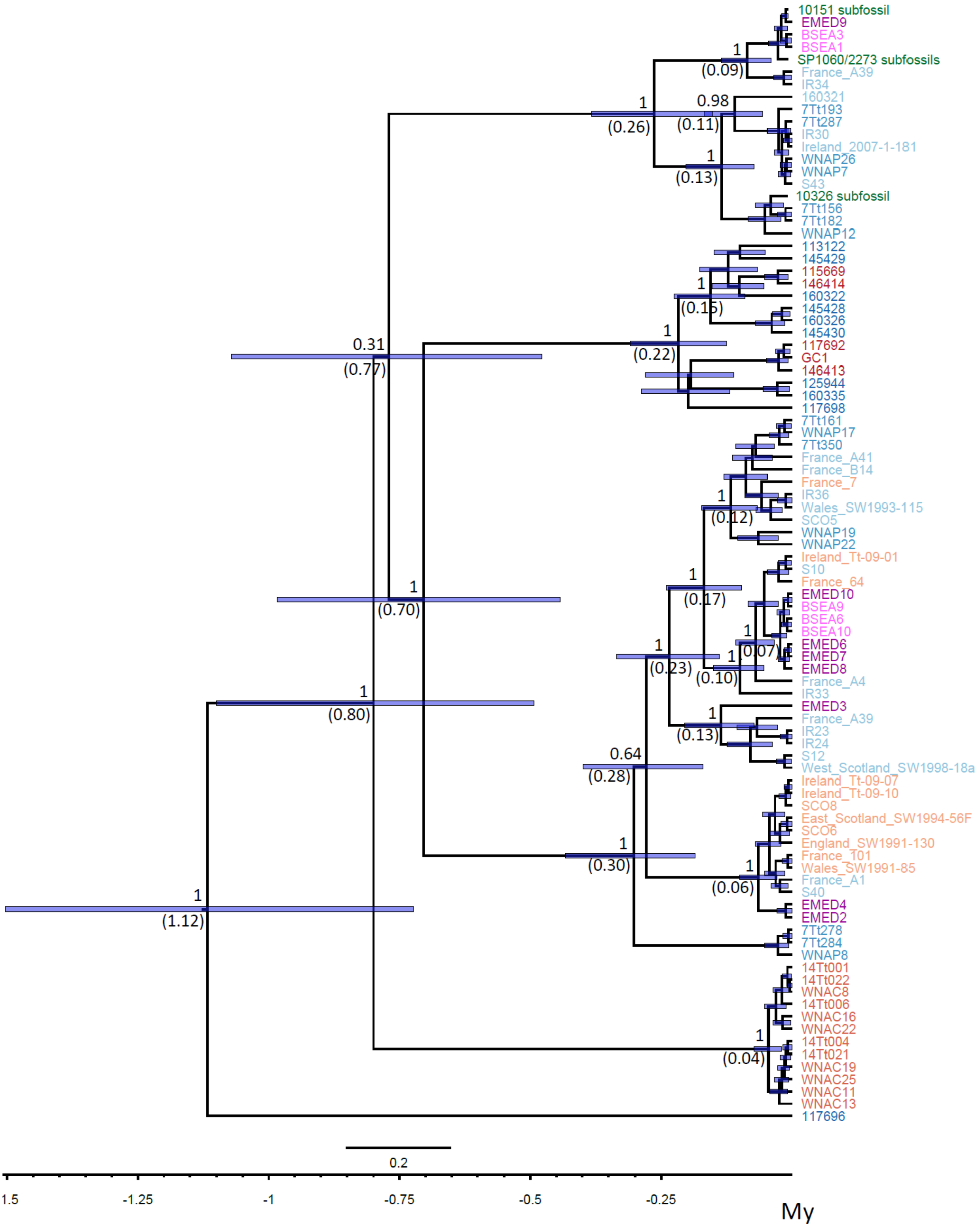

Figure S2. Time-calibrated phylogenetic tree for the *Tursiops truncatus* samples estimated with BEAST2 coalescent model with constant populations and using 13 mitochondrial coding genes. The numbers above and below nodes represent the node posterior probability and mean node age (in brackets), respectively, and the bars depict 95% HPDI in node TMRCA. The ancient subfossil samples are highlighted in green colour. The pelagic populations are coloured in different shades of blue, and the coastal populations of red. Note that the eastern Mediterranean Sea samples are in purple and the Black Sea in pink.

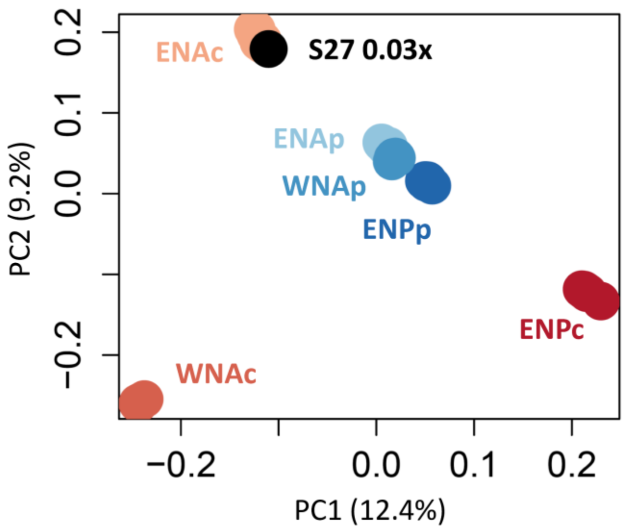

Figure S3. Principal component analysis of pseudo-haploid data from 55 modern and 1 projected downsampled sample (S27 from the ENAc), mapped to the killer whale reference genome, showing first and second principal components (PCs) based on 612,694 SNPs.

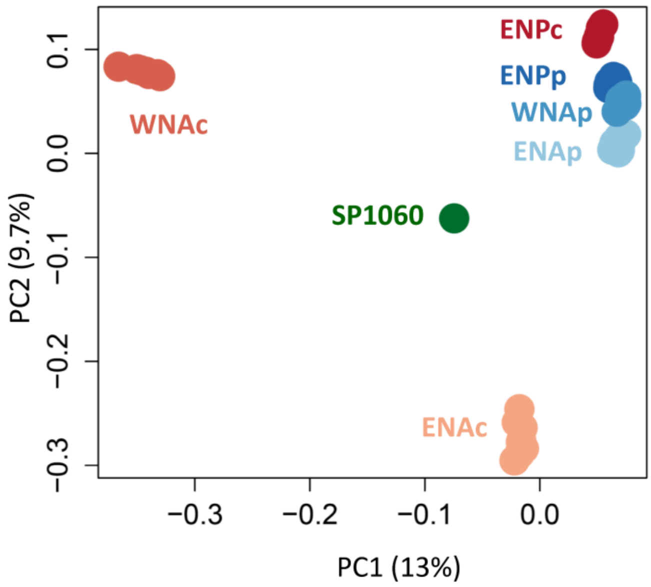

Figure S4. Principal component analysis (PCA) projections of SP1060, mapped to the bottlenose dolphin reference genome using BWA relaxed parameters for ancient DNA, on the principal components segregating the modern genome with smartpca using call genotypes and including transversions, sites with individual minimum coverage of 3x and no missing data in SP1060 for a total of 112,706 SNPs. First and second components are shown.

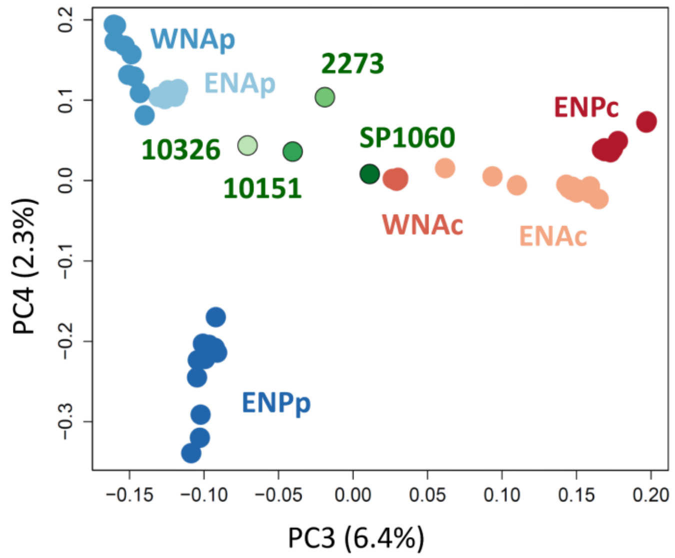

Figure S5. Principal component analysis (PCA) projections of the four ancient samples, mapped to the killer whale reference genome, on the principal components segregating the modern genomes using pseudo-haploid data and removing transversions for a total of 624,969 SNPs. Third and fourth components are shown.

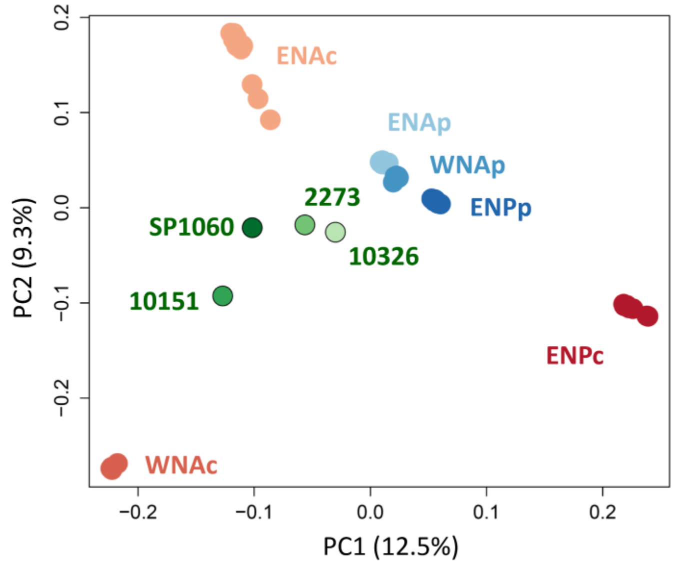

Figure S6. Principal component analysis (PCA) projections of the four ancient samples, mapped to the bottlenose dolphin reference genome using BWA relaxed parameters for ancient DNA, on the principal components segregating the modern genomes using pseudo-haploid data and removing transversions for a total of 885,285 SNPs. First and second components are shown.

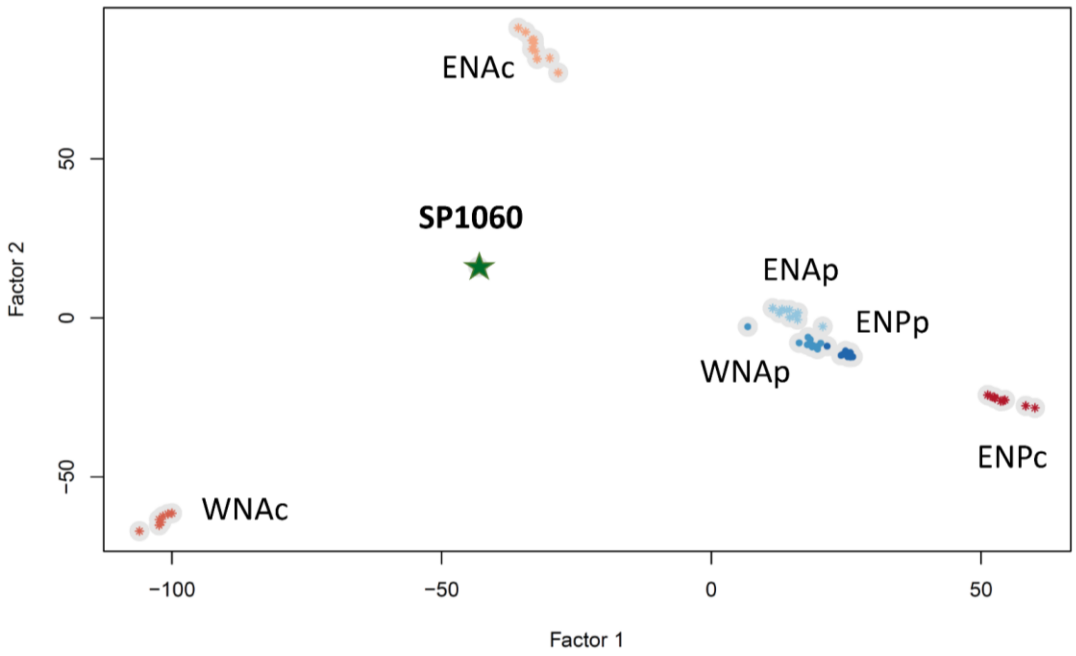

Figure S7. Factor analysis of 57 modern common bottlenose dolphins and one ancient common bottlenose dolphin of age 5,979-5,626 years BP. 112,506 SNPs with no missing data in SP1060 were included, missing genotypes were imputed in the modern individuals. Data was mapped to the bottlenose dolphin reference genome with relaxed parameters.

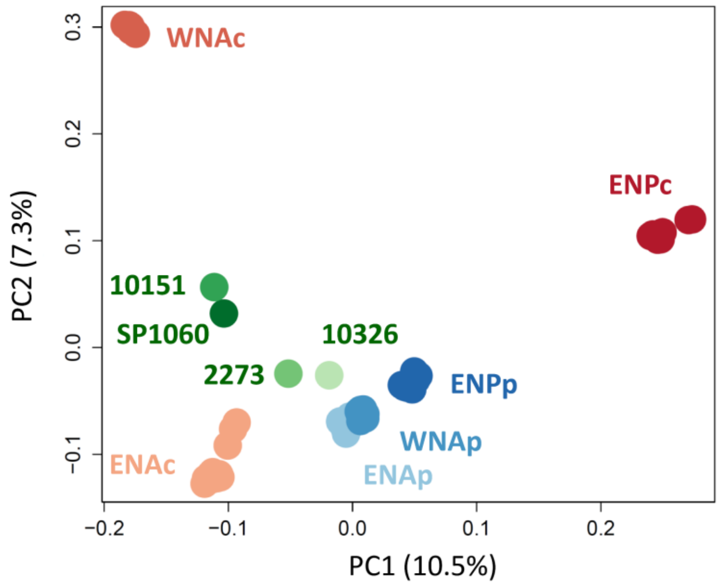

Figure S8. Principal components analyses using the single read sampling in ANGSD showing the first and second components for the 60 modern samples, SP1060, NMR2273, NMR10326 and NMR10151 mapped to the bottlenose dolphin reference genome with relaxed parameters, removing transitions, for a total of 16,286 SNPs.

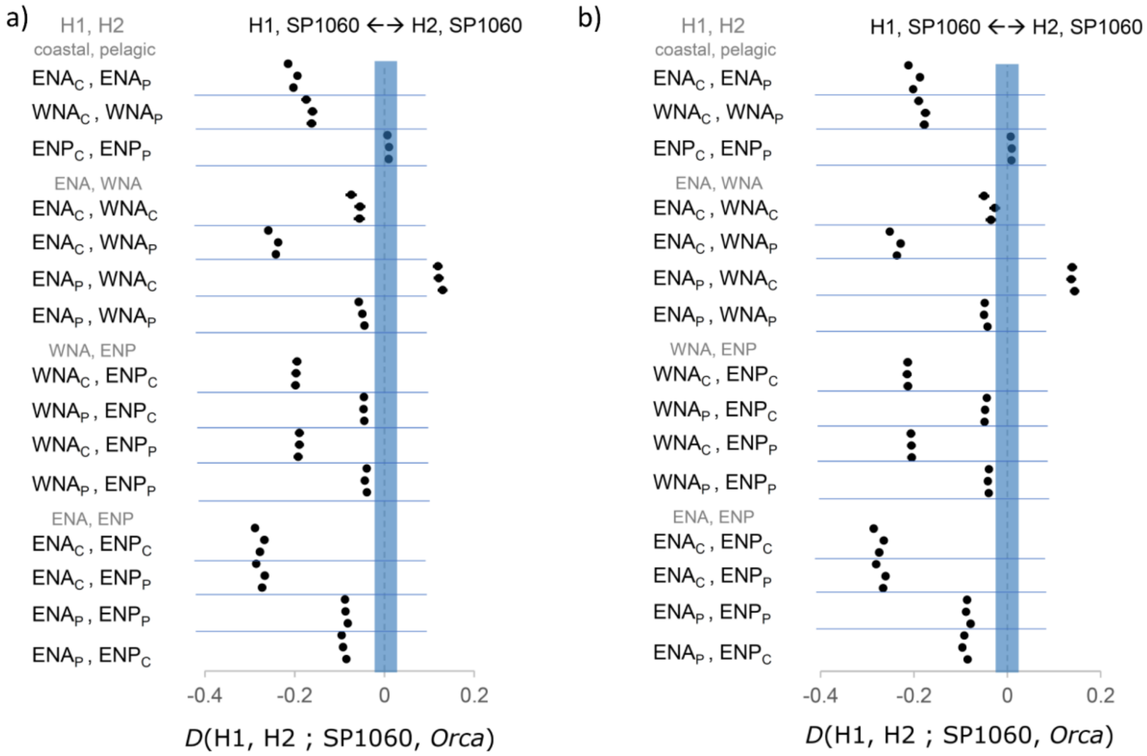

Figure S9. Plots of the D-statistic (ABBA-BABA) test *D*(H1,H2; SP1060, *Orca*) for the dataset mapped to the bottlenose dolphin reference genome with relaxed parameters a) without the transitions and b) with the transitions. All possible 15 combinations of two modern dolphin populations were included as the in-group (H1 and H2). For each combination of in-groups, three comparisons with different individuals were computed. Blue shading indicates non-significant results, $-3 > Z < 3$. SP1060 did not share any significant excess of derived alleles (non-significant *D* of 0.010-0.12) with either the ENPc and ENPp, further confirming independent evolution of coastal populations in the Atlantic and the Pacific (see Louis et al. 2021 [6]).

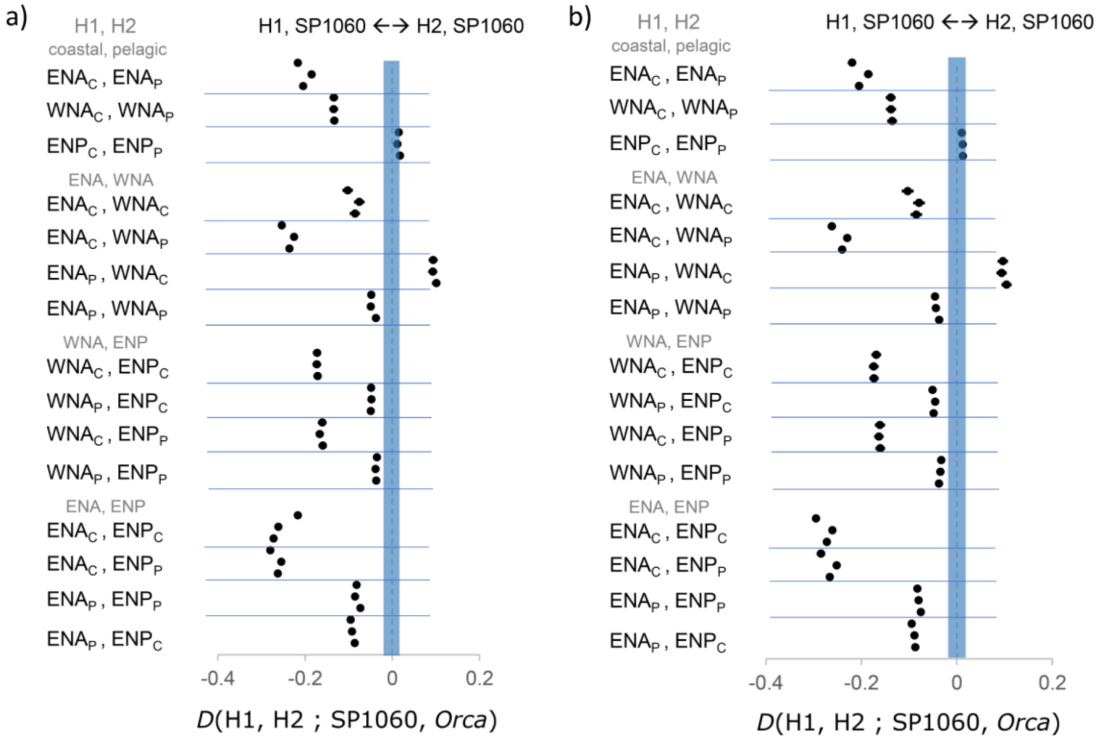

Figure S10. Plots of the D-statistic (ABBA-BABA) test *D*(H1,H2; SP1060, *Orca*) for the dataset mapped to the killer whale reference genome a) without the transitions and b) with the transitions. All possible 15 combinations of two modern dolphin populations were included as the in-group (H1 and H2). For each combination of in-groups, three comparisons with different individuals were computed. Blue shading indicates non-significant results, $-3 > Z < 3$.

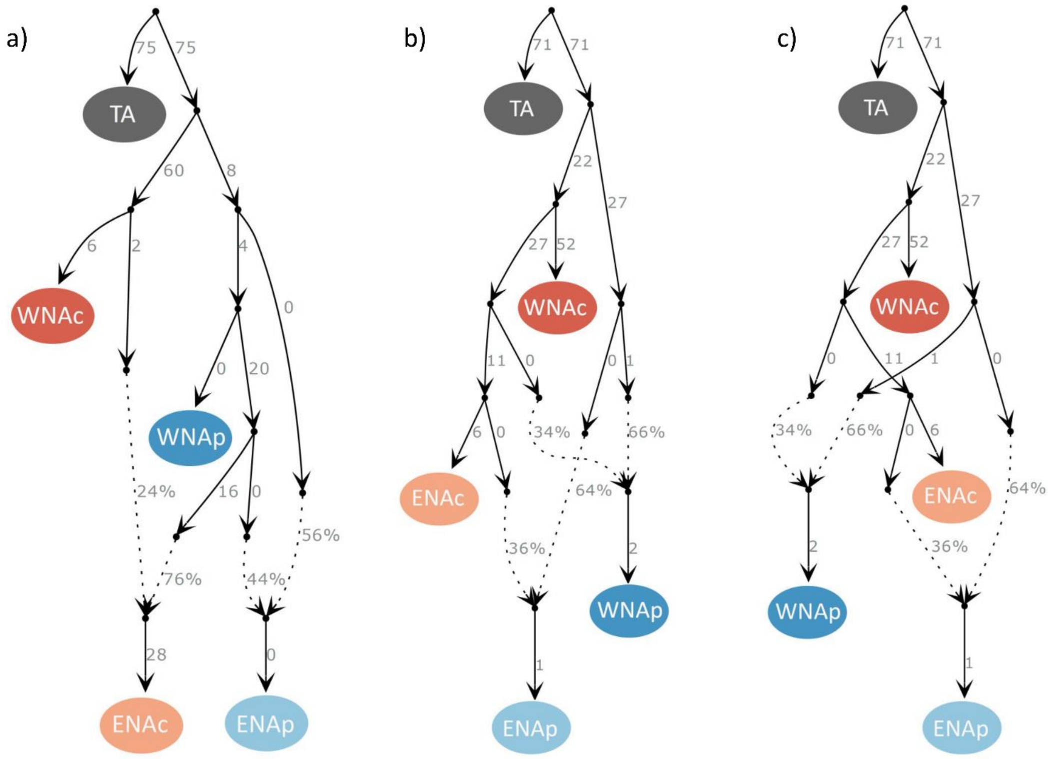

Figure S11. Evolutionary relationships between the North Atlantic modern bottlenose dolphin populations. Populations include eastern North Atlantic coastal (ENAc) and pelagic (ENAp) populations, and western North Atlantic coastal (WNAc) and pelagic (WNAp) populations, and the outgroup is the Indo-Pacific bottlenose dolphin (*Tursiops Aduncus*, TA). Admixture graphs were built using called genotypes with data mapped to the bottlenose dolphin reference genome including 223,950 SNPs. Continuous lines indicate phylogenetic relationships between populations/samples and the numbers at their right side the estimated genetic drift. Dotted lines show admixture edges and the number at their right side the percentage of ancestry deriving from each lineage.

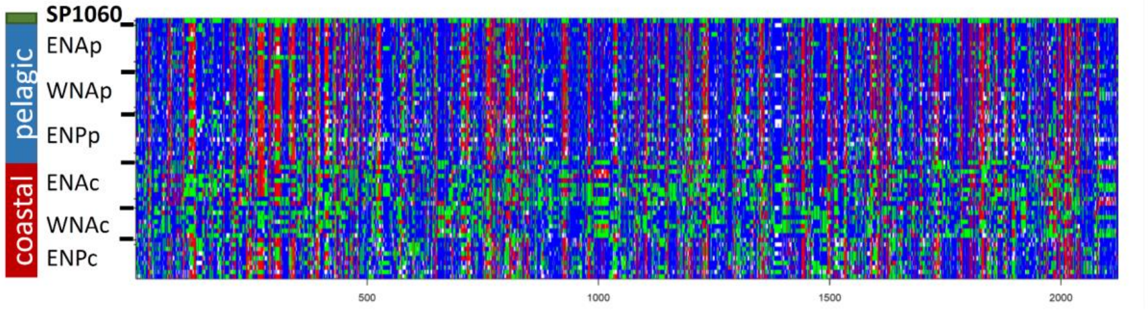

Figure S12. Genotypes of the SNPs under repeated selection to coastal habitat in modern individuals and ancient individual SP1060. These SNPs include 2,122 SNPs with no missing data in SP1060 out of the 7,165 SNPs identified in Louis et al. 2021. Plot of the homozygote reference genotypes in blue, heterozygote in green and homozygote derived in red.

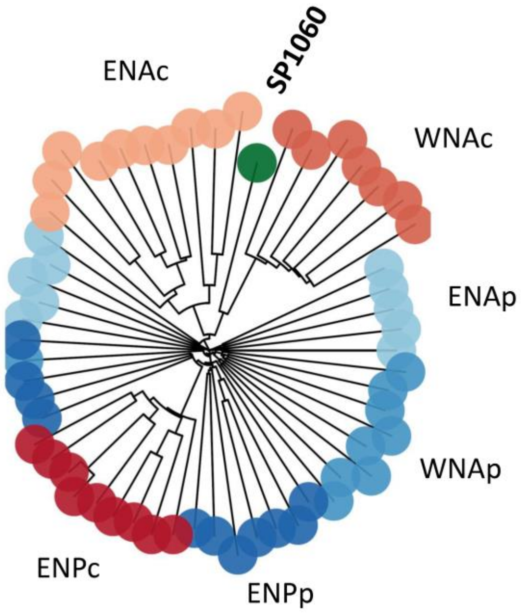

Fig S13. Patterns of genetic variation of 2,122 neutral SNPs in modern individuals and ancient individual SP1060. Neighbour-joining distance tree showing the genetic structure of the common bottlenose dolphin samples for this particular SNP set, which show no missing data in SP1060.

Table S1. Ancient subfossil specimen collection number, site location, radio-carbon dating reference code, radiocarbon-dating age (year BP), recalibrated age (year BP) for 1 sigma (1σ, 68% CI) and 2 sigma (2σ, 95% CI).

| Sample | Site | Code | $^{14}$C Age (BP) | Calibrated $^{14}$C age (BP) [1σ | Calibrated $^{14}$C age (BP) [2σ] |
|---|---|---|---|---|---|
| **SP1060** | Southern Bight | R-EVA 1656 | 5,624 +/- 28 | 5,896-5,723 | 5,979-5,626 |
| **NMR10326** | Smiths Knoll | GrA 25851 | 8,135 +/- 45 | 8,518-8,346 | 8,610-8,243 |
| **NMR 2273** | Smiths Knoll | GrA 25850 | 7,390 +/- 50 | 7,745 -7,572 | 7,842-7,492 |
| **NMR10151** | Southern Bight, 52:53:842N 2:83:471E | OxA-32156 | 6,822 +/- 39 | 7,228-7,036 | 7,303-6,935 |

Table S2. Summary statistics for the nuclear data for a) the ancient samples and b) the three new modern samples. “Reference” indicates the reference genome used for mapping, “sample” indicate sample name, “nb” the number of libraries prepared for that samples, “total reads” the raw number of reads, “mapped N” the number of reads mapped to the nuclear reference genome, “mapped N q25 unique” the number of reads mapped to the nuclear reference genome after mapping quality filter of 25 and removing duplicates, “%N” is the endogenous content after mapping quality and duplicate filters, “effective coverage” represents the coverage after all filters (i.e. post-QC filtering, repeat masking, removing duplicates, base quality recalibration, regions of excessive coverage, sex chromosomes and scaffolds shorter than 10 Mbp, see details in Louis et al. 2021 [6]).

a)

| **reference** | **sample** | **nb** | **total reads** | **mapped N** | **mapped N q25 unique** | **% N** | **effective coverage** |
|---|---|---|---|---|---|---|---|
| *T. truncatus* | SP1060 | 4 | 527,270,096 | 225,493,587 | 147,583,308 | 27.99 | 2.841 |
| *O. orca* | SP1060 | 4 | 527,270,096 | 191,133,611 | 127,139,634 | 24.11 | 2.976 |
| *T. truncatus* | NMR2273 | 5 | 87,203,486 | 57,760,143 | 744,073 | 0.85 | 0.008 |
| *O. orca* | NMR2273 | 5 | 87,203,486 | 46,032,421 | 544,427 | 0.62 | 0.001 |
| *T. truncatus* | NMR10326 | 9 | 113,642,049 | 97,736,589 | 1,658,557 | 1.46 | 0.023 |
| *O. orca* | NMR10326 | 9 | 113,642,049 | 79,607,214 | 1,254,302 | 1.10 | 0.002 |
| *T. truncatus* | NMR10151 | 7 | 83,440,588 | 50,960,537 | 102,666 | 0.12 | 0.001 |
| *O. orca* | NMR10151 | 7 | 83,440,588 | 43,138,459 | 99,870 | 0.12 | 0.001 |

b)

| **reference** | **sample** | **nb** | **total reads** | **mapped N q25 unique** | **effective coverage** |
|---|---|---|---|---|---|
| ***T. truncatus*** | Normandy | 1 | 75,898,164 | 59,925,176 | 1.732 |
| ***O. orca*** | Normandy | 1 | 75,898,164 | 59,892,951 | 1.745 |
| ***T. truncatus*** | West Scotland | 1 | 156,592,102 | 119,308,303 | 3.432 |
| ***O. orca*** | West Scotland | 1 | 156,592,102 | 118,514,741 | 3.439 |
| ***T. truncatus*** | Shannon | 1 | 97,573,706 | 74,098,872 | 2.912 |
| ***O. orca*** | Shannon | 1 | 97,573,706 | 73,958,864 | 2.218 |

Table S3. Whole mtDNA sequences/mitochondrial haplotypes downloaded from GenBank and used in the phylogenetic and coalescent analyses. Only the protein coding regions from these sequences were used in the estimation of time-calibrated phylogenies for delphinids and for *T. truncatus*.

| **Delphinid phylogeny** | | |
|---|---|---|
| **Species** | **Accession number** | **GenBank reference** |
| *Cephalorhynchus heavisidii* | JN632624 | Hassanin *et al.*, 2012 [29] |
| *Orcaella brevirostris* | JF289177 | Vilstrup *et al.*, 2011 [30] |
| *Orcaella heinsohni* | JF339977 | Vilstrup *et al.*, 2011 |
| *Peponocephala electra* | JF289175 | Vilstrup *et al.*, 2011 |
| *Feresa attenuata* | JF289171 | Vilstrup *et al.*, 2011 |
| *Globicephala melas* | JF339972 | Vilstrup *et al.*, 2011 |
| *Globicephala macrorhynchus* | JF339976 | Vilstrup *et al.*, 2011 |
| *Pseudorca crassidens* | JF289173 | Vilstrup *et al.*, 2011 |
| *Grampus griseus* | EU557095 | Xiong *et al.*, 2009 [31] |
| *Stenella attenuata* | EU557096 | Xiong *et al.*, 2009 |
| *Stenella coeruleoalba* | EU557097 | Xiong *et al.*, 2009 |
| *Delphinus capensis* | EU557094 | Xiong *et al.*, 2009 |
| *Sousa chinensis* | EU557091 | Xiong *et al.*, 2009 |
| *Lagenorhynchus albirostris* | NC005278 | Arnason *et al.*, 2004 [32] |
| *Orcinus orca*, resident ecotype | GU187192 | Morin *et al.*, 2010 [45] |
| *Orcinus orca*, transient ecotype | GU187173 | Morin *et al.*, 2010 |
| *Steno bredanensis* | JF339982 | Vilstrup *et al.*, 2011 |
| *Tursiops aduncus* | KF570335 | Moura *et al.*, 2013 [16] |
| *Tursiops australis* | KF570363 | Moura *et al.*, 2013 |

| ***Tursiops truncatus* phylogeny** | | | |
|---|---|---|---|
| **Species** | **Haplotype name** | **Accession number** | **GenBank reference** |
| *T. truncatus* | EMED3 | KF570315 | Moura *et al.*, 2013 |
| *T. truncatus* | EMED4 | KF570316 | Moura *et al.*, 2013 |
| *T. truncatus* | EMED5 | KF570317 | Moura *et al.*, 2013 |
| *T. truncatus* | EMED1 | KF570318 | Moura *et al.*, 2013 |
| *T. truncatus* | EMED2 | KF570319 | Moura *et al.*, 2013 |
| *T. truncatus* | EMED10 | KF570320 | Moura *et al.*, 2013 |
| *T. truncatus* | EMED6 | KF570321 | Moura *et al.*, 2013 |
| *T. truncatus* | EMED9 | KF570322 | Moura *et al.*, 2013 |
| *T. truncatus* | EMED7 | KF570323 | Moura *et al.*, 2013 |
| *T. truncatus* | EMED8 | KF570324 | Moura *et al.*, 2013 |
| *T. truncatus ponticus* | BSEA2 | KF570325 | Moura *et al.*, 2013 |
| *T. truncatus ponticus* | BSEA3 | KF570326 | Moura *et al.*, 2013 |
| *T. truncatus ponticus* | BSEA1 | KF570327 | Moura *et al.*, 2013 |
| *T. truncatus ponticus* | BSEA6 | KF570328 | Moura *et al.*, 2013 |
| *T. truncatus ponticus* | BSEA7 | KF570329 | Moura *et al.*, 2013 |

| | | | |
|---|---|---|---|
| *T. truncatus ponticus* | BSEA5 | KF570330 | Moura *et al.*, 2013 |
| *T. truncatus ponticus* | BSEA4 | KF570331 | Moura *et al.*, 2013 |
| *T. truncatus ponticus* | BSEA8 | KF570332 | Moura *et al.*, 2013 |

Table S4. Best partitioning schemes for nucleotide substitution models used in the construction of the delphinid and *T. truncatus* time-calibrated phylogenies.

| **Delphinids: All codon model** | | |
|---|---|---|
| **Partition** | **Best model** | **Subset partitions** |
| p1 | HKY+I+G | atp6, atp8, cox2, cox3, cytb, nd1, nd2, nd3, nd4, nd4l, nd5 |
| p2 | HKY+I+G | cox1 |
| p3 | HKY+I+G | nd6 |
| Delphinids: Third codon only model | | |
| **Partition** | **Best model** | **Subset partitions** |
| p1 | TrN+I+G | atp6, atp8, cox3, cytb, nd1, nd2, nd3, nd4, nd4l, nd5 |
| p2 | TrN+G | atp8, cox1 |
| p3 | TrN+G | nd6 |
| *T. truncatus*: All codon model | | |
| **Partition** | **Best model** | **Subset partitions** |
| p1 | HKY+I | atp6, atp8, cox3, cytb, nd1, nd2, nd3, nd4, nd4l, nd5 |
| p2 | HKY+G | cox1, cox2 |
| p3 | HKY+I | nd6 |
| *T. truncatus*: Third codon only model | | |
| **Partition** | **Best model** | **Subset partitions** |
| p1 | TrN | cox3, cytb, nd1, nd2, nd4, nd4l, nd5 |
| p2 | TrN | atp6, atp8, cox1, cox2, nd3 |
| p3 | TrN+I | nd6 |

Table S5. Different coalescent scenarios run with BEAST2 to estimate *T. truncatus* phylogeny.

| Tree scenario | Model | Root calibration, mean (SD) | Tip calibration, mean (SD) | | |
|---|---|---|---|---|---|
| | | | **2273** | **3920** | **10151** |
| Without 117696, without 10151 | All codon | 0.9162 (0.158) | 0.007663 (1.00E-04) | 0.00843 (1.00E-04) | NA |
| Without 117696, without 10151 | 3rd codon | 0.7767 (0.145) | 0.007663 (1.00E-04) | 0.00843 (1.00E-04) | NA |
| Without 117696, with 10151 | All codon | 0.9162 (0.158) | 0.007663 (1.00E-04) | 0.00843 (1.00E-04) | 0.007126 (1.00E-04) |
| Without 117696, with 10151 | 3rd codon | 0.7767 (0.145) | 0.007663 (1.00E-04) | 0.00843 (1.00E-04) | 0.007126 (1.00E-04) |
| With 117696, without 10151 | All codon | 1.174 (0.2) | 0.007663 (1.00E-04) | 0.00843 (1.00E-04) | NA |
| With 117696, without 10151 | 3rd codon | 1.0332 (0.18) | 0.007663 (1.00E-04) | 0.00843 (1.00E-04) | NA |
| With 117696 and 10151 | All codon | 1.174 (0.2) | 0.007663 (1.00E-04) | 0.00843 (1.00E-04) | 0.007126 (1.00E-04) |
| With 117696 and 10151 | 3rd codon | 1.0332 (0.18) | 0.007663 (1.00E-04) | 0.00843 (1.00E-04) | 0.007126 (1.00E-04) |
